# The effect of noise trauma and high-frequency stimulation on thalamic sensory gating in rodents

**DOI:** 10.1101/2020.04.03.023143

**Authors:** Aryo Zare, Gusta van Zwieten, Sonja A. Kotz, Yasin Temel, Benjamin G. Schultz, Michael Schwartze, Marcus L.F. Janssen

## Abstract

**Background:** The medial geniculate body (MGB) of the thalamus plays a central role in tinnitus pathophysiology. Breakdown of sensory gating in this part of the auditory thalamus is a potential mechanism underlying tinnitus. The alleviation of tinnitus-like behavior by high-frequency stimulation (HFS) of the MGB might mitigate dysfunctional sensory gating.

**Objective:** The study aims at exploring the role of the MGB in sensory gating as a mandatory relay area in auditory processing in noise-exposed and control subjects, and to assess the effect of MGB HFS on this function.

**Methods:** Noise-exposed rats and controls were tested. Continuous auditory sequences were presented to allow assessment of sensory gating effects associated with pitch, binary grouping, and temporal regularity. Evoked potentials (EP) were recorded from the MGB and acquired before and after HFS (100 Hz).

**Results:** Noise-exposed rats showed differential modulation of MGB EP amplitudes, confirmed by significant main effects of stimulus type, pair position and temporal regularity. Noise-exposure selectively abolished the effect of temporal regularity on EP amplitudes. A significant three-way interaction between HFS phase, temporal regularity and rat condition (noise-exposed, control) revealed that only noise-exposed rats showed significantly reduced EP amplitudes following MGB HFS.

**Conclusion:** This is the first report that shows thalamic filtering of incoming auditory signals based on different sound features. Noise-exposed rats further showed higher EP amplitudes in most conditions and did not differentiate the temporal regularity. Critically, MGB HFS was effective in reducing amplitudes of the EP responses in noise-exposed animals.

**Highlights:**

- EP findings indicate sensory gating in the MGB in rats.
- Noise exposure alters EP amplitudes in the MGB.
- HFS selectively suppresses EP responses in noise-exposed animals.

## Introduction

Tinnitus sufferers perceive a sound in the absence of an external audible sound source. Patients suffering from tinnitus often experience a substantially lower quality of life and effective therapies are still lacking. A limited understanding of tinnitus pathophysiology hinders the development of effective treatments. Although the specific pathophysiological mechanisms underlying tinnitus remain elusive, multiple theories with overlapping neurophysiological mechanisms have been proposed. The central gain model proposes that tinnitus results from dynamic compensatory gain enhancement [1]. Accordingly, there is an increase of evoked neural responses following noise trauma [2]. More specifically, increased ‘gain’ in auditory neurons would lead to alterations in synaptic transmission as well as neuronal excitability and synchrony [2, 3]. Furthermore, breakdown of sensory gating (SG) has been proposed as a mechanism underlying phantom auditory perception [5, 6]. All mechanisms might contribute to altered oscillations within the thalamo-cortical loop, described as thalamo-cortical dysrhythmia [4].

SG is defined as the adaptive filtering of changing stimulus features (gating in) relative to repetitive stimulus features (gating out) [5, 6]. According to this model, the tinnitus sensation is linked to the failure of sensory-perceptual filtering or “noise cancelling system”. The medial geniculate body (MGB) of the thalamus is a major relay between midbrain and cortex, and acts as the gatekeeper for auditory signals. Therefore, we expected that the MGB plays an important role in dysfunctional SG in tinnitus. How this thalamic gate functions and how it can potentially be modulated, remains poorly understood [7]. It has been hypothesized that the MGB is mainly inhibited by the thalamic reticular nucleus, a structure that derives input from a number of paralimbic structures [8]. This inhibitory feedback loop may block irrelevant sensory inputs and this mechanism might be altered in tinnitus. To our knowledge, only cortical features of SG in tinnitus have been directly studied so far [9, 10].

SG may operate on the basis of different features of auditory input, including the specific stimulus type and timing, (i.e. sound structure and temporal arrangement in terms of grouping and regularity). Together, these features define the parameters that guide filtering. Such filtering dismisses goal-irrelevant neural signals in favor of selective attention to goal-relevant information [11]. Dysfunctional SG is evident in various neuropsychiatric conditions, such as impulsivity control disorders and schizophrenia [12, 13]. Recent evidence points to the possibility of tinnitus as a possible further condition in which impaired SG plays a critical role. Correlations between tinnitus severity and decreased SG via the Pa component of cortical auditory evoked potentials and an increased SG via the N1 component have recently been described [9]. SG is also influenced by state changes of the organism [14]. For example, a case-control study showed that there is a relation between the level of SG and behavioral aspects of tinnitus based on the tinnitus handicap and sensory gating inventories [10]. However, it is unclear how SG is implemented in thalamic processing stages, and if SG can be modulated in the specific context of tinnitus.

Deep brain stimulation (DBS) is a potential method of exerting a modulatory influence on auditory SG in tinnitus. DBS has recently emerged as a promising treatment option for tinnitus [15–17]. High-frequency stimulation (HFS) applied to the auditory pathway has been shown to effectively reduce tinnitus-like behavior in rats [18–20]. The exact working mechanisms of DBS remain elusive, although complex inhibitory and distant excitatory effects have been described [15, 21].

In the current study, we tested the hypothesis that SG occurs at the level of the MGB and can be altered in a noise-exposed tinnitus animal model. Considering the positive effect of MGB HFS on tinnitus-like behavior, we hypothesize that HFS of the MGB modulates thalamic SG. In other words, HFS might restore a dysfunctional “noise cancelling system” by counteracting aberrant filtering of sensory input at the level of the MGB [21].

To this end, we investigated auditory SG in the MGB in control and noise-exposed tinnitus rats and further explored the effect of HFS. Tinnitus was assessed using gap-prepulse inhibition of acoustic startle (GPIAS). EP recordings were conducted in anesthetized animals, using paired-tone auditory stimulus sequences that were composed of different pitches (frequent standard and infrequent deviant tones), position (temporal order within the pair) and temporal regularity (regular [isochronous] and irregular), before and after MGB HFS.

## Material and Methods

### Subjects

The experimental group used for the current study has been described before [van Zwieten et al, *under review*]. Briefly, a total of 13 male Sprague Dawley rats were used, divided into two groups: i) noise-exposed tinnitus (n=8) and ii) unexposed controls (n=5). Rats were individually housed in standard Makrolon™ cages, with food and water *ad libitum*. Conditions in the room were constant, with a temperature of 20°C to 22°C and humidity of 60% to 70%. The light-dark cycle was reversed, and experiments were conducted within the dark period. The study was approved by the Animal Experiments and Ethics Committee at Maastricht University, the Netherlands.

### Study Design

The overall design of the study comprised four main parts: GPIAS, noise exposure, repeated GPIAS, and recording of EPs and HFS. The following readout parameters were used: the dependent variable was the amplitude of the evoked potential (mV) in response to the stimulus onset. The independent variables were noise exposure (noise-exposed, unexposed; between-subjects), tone position (first, second; within-subjects), timing (regular, irregular; within-subjects), and HFS phase (pre-HFS, post-HFS; within-subjects). Tinnitus induction, GPIAS, surgical and deep brain stimulation procedures have been described in detail elsewhere [van Zwieten et al, *under review*].

### Tinnitus Induction

Animals were anesthetized using Ketamine and Xylazine. Rats in the noise-exposed group (i) were unilaterally exposed to a 16 kHz octave-band noise at 115 dB for 90 min. Unexposed control rats (ii) were only anesthetized.

### Gap Prepulse Inhibition of the Acoustic Startle

Behavioral evidence of tinnitus was assessed before and three weeks after noise or sham exposure by GPIAS, previously described in detail [van Zwieten et al, *under review*]. Freely moving rats were placed in a cylinder, on a piezo sensor. Gap and no-gap trials were alternately executed with 20 repetitions per background frequency (10 kHz, 16 kHz and 20 kHz at 75 dB). The startle stimulus consisted of a click sound of 105 dB intensity with 20 ms duration. In gap trials, a gap of 50 ms was added to the background sound, 100 ms prior to the startle stimulus. Prior to each session, the animals were acclimatized for 5 minutes and habituated to the startle sound by presenting 10 no-gap trials. To habituate the animals to the testing procedure, one complete session was performed at the beginning of the experiment. Two complete sessions per condition were conducted, each on separate testing days. The gap/no-gap ratio was calculated by dividing the amplitude per gap-startle with the mean of all startle-only trials [19].

### Surgical procedure

Rats were anesthetized by intraperitoneal administration of urethane (7.5 ml/kg loading dose and 0.3 ml repetitive dose for maintenance) from a 20% of weight urethane solution (Sigma-Aldrich / Merck KGaA, Darmstadt, Germany). The level of anaesthesia was monitored by checking whisker and pedal reflexes. Body temperature was controlled and maintained at 37°C by means of a heating pad (ATC1000, World Precision Instruments, Sarasota, Florida). Rats were mounted on a stereotaxic apparatus (Stoelting Co, Illinois, USA) using hollow ear bars to allow presentation of auditory stimuli. A craniotomy was performed in order to access the left MGB which is contralateral to the noise (or sham) exposed ear. To record local field activity, a bipolar electrode was inserted into the contralateral MGB (craniocaudal −5.5mm, mediolateral +3.6mm, dorsoventral −6mm) according to Paxinos rat brain atlas [22]. The electrode was a custom-made platinum-iridium bipolar electrode with a shaft diameter of 250 μm and tip diameter of 50 μm (Technomed, Beek, the Netherlands).

### Acquisition of evoked potentials and the auditory stimulation paradigm

Electrophysiological recordings were performed four weeks after noise exposure. The electrode was connected to a data acquisition system (PowerLab 8/35, New South Wales, Australia) and sampled at 20000 Hz to record the local field potentials, using LabChart Pro 7 software (ADInstruments, Castle Hill, Australia). A band pass filter of 0.1 Hz – 1 kHz was used. External auditory stimuli were generated by a PC audio interface (0204 USB Audio Interface, E-MU systems, Dublin, Ireland), using custom-made MATLAB scripts. Auditory stimuli were amplified and presented with an Ultrasonic Dynamic Speaker (Vifa, Avisoft Bioacoustics, Berlin, Germany), calibrated with a modular precision sound level meter (Bruel and Kjaer 2231, San Diego, USA) and a free-field microphone (Bruel and Kjaer type 4191, San Diego, USA). The contralateral ear was sealed to block external auditory perception.

The auditory stimuli consisted of two pure tones with different pitches, the 600 Hz tone serving as the standard and the 660 Hz tone as deviant (see Figure 1). The stimulus sequences consisted of six blocks of binary grouped tones, with each block lasting for 1 minute. Each block thus consisted of 60 stimulus pairs. Six blocks were performed without interruption for a total of six minutes, once with regular (isochronous) timing intervals and once with irregular (jittered) intervals. Occurrence of the deviant tones was balanced across the two positions of each pair. Each stimulus lasted for 70 ms including 10 ms rise and fall times. The standard-to-deviant ratio was 4:1. These stimuli were organized into two separate arrangements, one generating a regular (predictable) and one an irregular (less predictable) timing. The latter was realized via random variation of both the interval within (intra-chunk) and the interval between consecutive pairs (inter-chunk interval). The pairs consequently either consisted of two standard tones (S1S2), or standard and then deviant tone (S1D2), or deviant then standard tone (D1S2). Each rat received an initial auditory stimulation with the regular and irregular sequence, both before and directly after HFS. A counterbalanced design was used. This setup allowed assessment of SG effects associated with pitch (stimulus type: standard or deviant), binary grouping (position in a pair: first or second), and temporal regularity (timing: regular or irregular).

**Figure 1.**
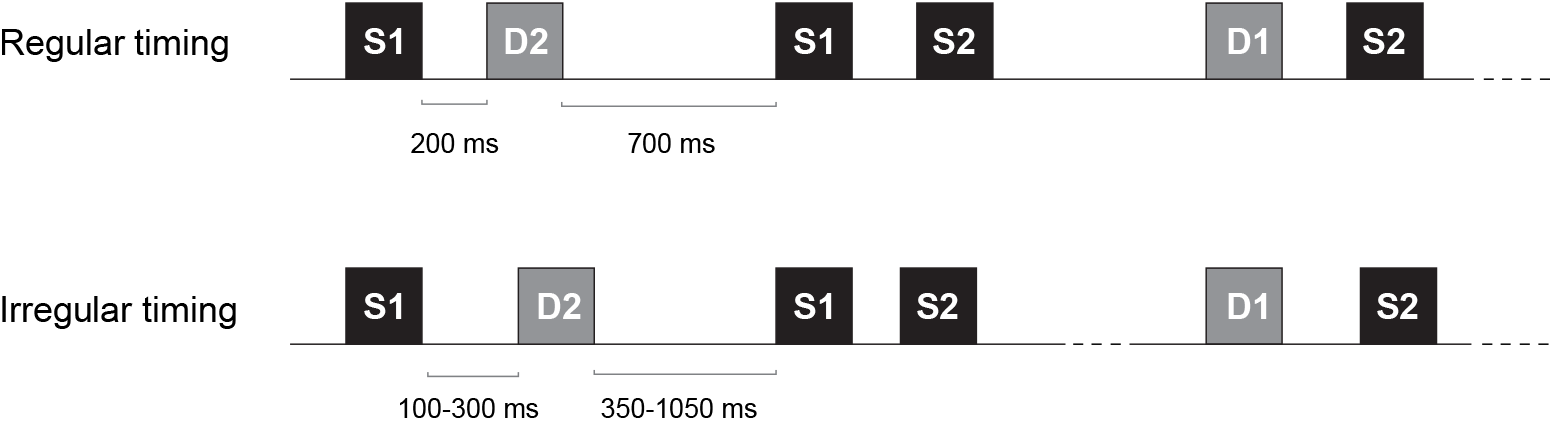
External auditory stimuli: An exemplary sequence of standard and deviant tones, depicting the length of intervals in regular and irregular timing conditions. Note that a range is designated in the irregular sequence. S = standard tone (600 Hz), D = deviant tone (660 Hz).

### Deep brain stimulation

MGB HFS was applied for a period of five minutes with the same bipolar electrode as used for the EP recordings. HFS (100 Hz, 60 μs, 100 μA, bipolar, monophasic square-wave pulses) was applied with a stimulator (DS8000, World Precision Instruments, Sarasota, Florida) connected to a constant-current isolator (DLS100, WPI, Sarasota, Florida). These stimulation parameters were based on previous experiments [20]. Regular and irregular external auditory sound sequences were repeated, each being preceded by HFS, again using a counterbalanced design.

### Electrode localization

To check for correct electrode tip placement, the rats were euthanized by decapitation while still being under general anesthesia, the brains were quickly removed and frozen in −40°C 2-methyl-butane (isopentane). The tissue was serially cut by cryostat (Leica CM3050S, Wetzlar, Germany). Hematoxylin-eosin staining was performed to confirm appropriate electrode placement.

### Data processing

Analyses were performed using the ‘Letswave’ toolbox [23] running in MATLAB®. The signals were bandpass filtered to include frequencies between 5-30 Hz. After baseline correction, outliers were removed for a maximum of 5 epochs (from a total of 72) in the deviant group and 20 in the standard group (from a total of 288). Next, data were averaged across each event code (S1, S2, D1, D2). Amplitude of the first peak following the trigger onset was defined for further statistical comparison (Figure 2).

**Figure 2.**
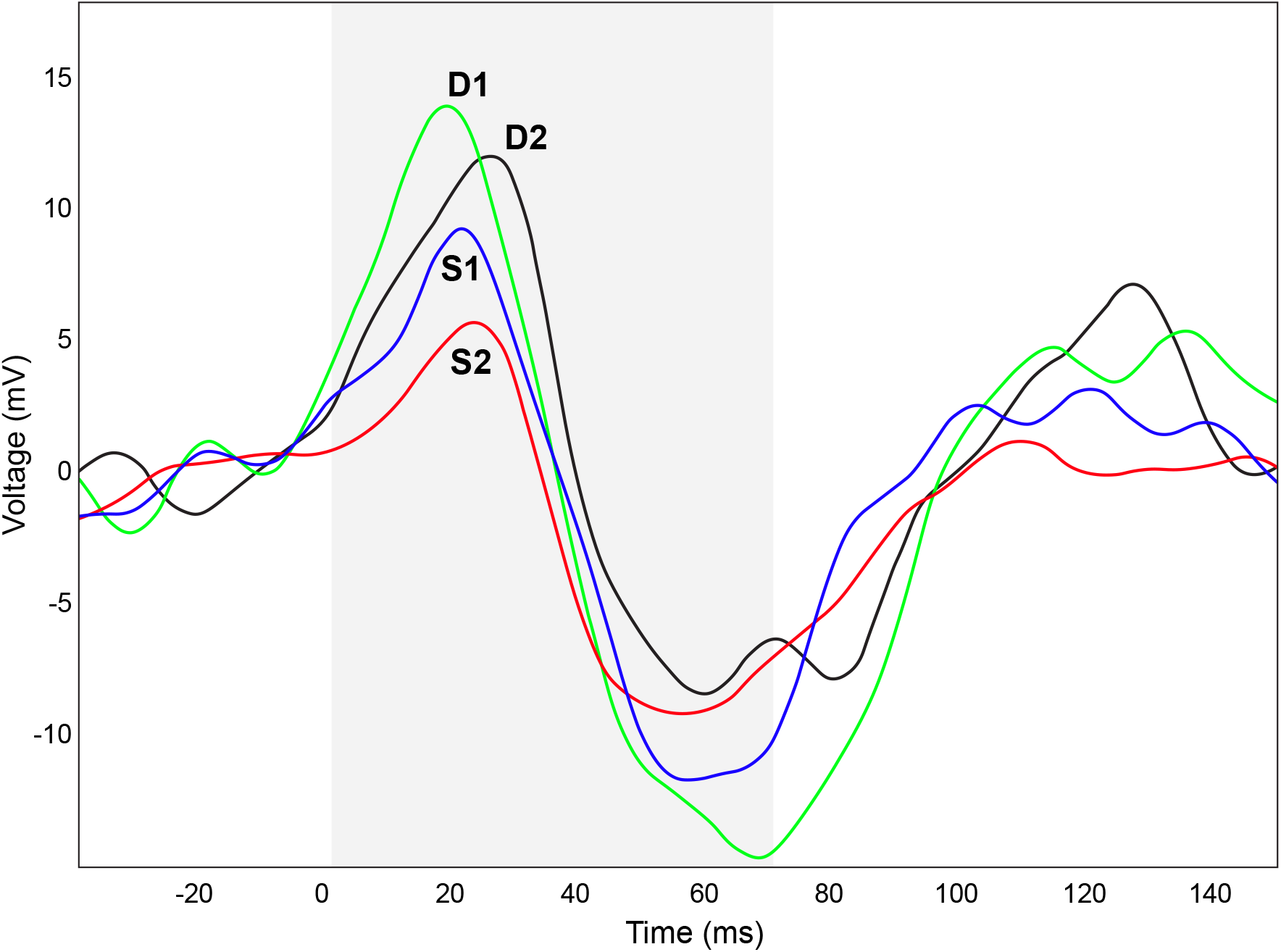
A representative mean evoked potential (EP) from a control rat at pre-HFS condition during the regular timing condition. Note that event codes (D1, D2, S1, S2) have different peak amplitudes. The grey area represents the auditory stimulus of 70 ms duration.

### Statistical analysis

Amplitudes were not normally distributed and, consequently, were log-transformed prior to analysis. Outliers were excluded based on visual inspection, specifically values less than −1 mV and greater than 1 mV. The log-transformed amplitudes were fitted to a *linear mixed-effects model (LMEM)* with fixed factors Stimulus Type (2; Standard, Deviant), Position (2; First, Second), Noise (2; Unexposed, Noise-exposed), Timing (2; Regular, Irregular), and HFS phase (2; Pre-HFS, Post-HFS), and random effects RatID, Block, and Trial, where Trial was nested in Block, and Block was nested in RatID. Models were selected based on the lowest *Akaike’s Information Criteria (AIC)* value when including the interaction term or only main effects between the fixed factors. Data were analyzed using the *lmer* function in the *lme4* package [24] for R software [25]. Effect sizes were measured as generalized *eta squared* (*η_G2_*) using the *aov_car* function in the *afex* package [26]. Pairwise comparisons were calculated as *Tukey’s Honestly Significant Difference (HSD)* using the *multcomp* package [27]. To determine evidence for the null hypothesis, *Bayes Factor t*-tests were performed using the *ttestBF* function of the *BayesFactor* package [28]. Following Jeffreys [29], *Bayes Factor* values were interpreted as follows: values around 1 indicates no evidence, 1-3 indicate anecdotal evidence, 3-10 indicate moderate evidence, 10-30 indicate strong evidence, 30-100 indicate very strong evidence, and greater than 100 indicates extreme evidence.

## Results

### Gap Prepulse Inhibition of Acoustic Startles

Tinnitus was assessed through GPIAS where the gap/no-gap ratios increased following noise exposure for 20 kHz and 16 kHz (*ps*<0.01) but not 10 kHz (*p*=0.21) background sound; unexposed rats showed no significant changes (*ps*>0.14). These findings suggest that tinnitus was successfully induced in the noise-exposed animals (van Zwieten et al. *under review*). These GPIAS results are similar to earlier published studies from our group [18, 19].

### Evoked potentials

Peak EP amplitude up to 70 ms after the trigger onset was measured to perform comparisons. The *LMEM* with the best model fit contained main effects of Stimulus Type and Position, and interactions between the other variables. There were significant main effects of Stimulus Type [*F* (1, 37256) = 4.20, *p*=0.04), *η_G2_*=0.001; Figure 3A], Position [*F* (1, 37255) = 10.86, *p*<0.001; Figure 3B], *η_G2_*= 0.001], and Timing [*F* (1, 37256) = 12.79, *p*<0.001), *η_G2_*=0.002]. There was a significant two-way interaction between Noise and Timing [*F* (1, 37256) = 21.83, *p*<0.001), *η_G2_*=0.003], and a significant three-way interaction between Noise, Timing, and HFS phase [*F* (1, 37256) = 12.67, *p*<0.001), *η_G2_* = .002]. No other main effects or interactions reached significance (*ps*>0.21). As shown in Figure 3A, deviant tones elicited significantly higher amplitudes than standard tones supporting the hypothesis that habituation occurs for stimuli that are more predictable and occur more often. Similarly, amplitudes were lower for the second stimulus in a pair compared to the first suggesting that sensory gating occurs for the second stimulus in a pair of temporally proximal tones (Figure 3B).

**Figure 3.**
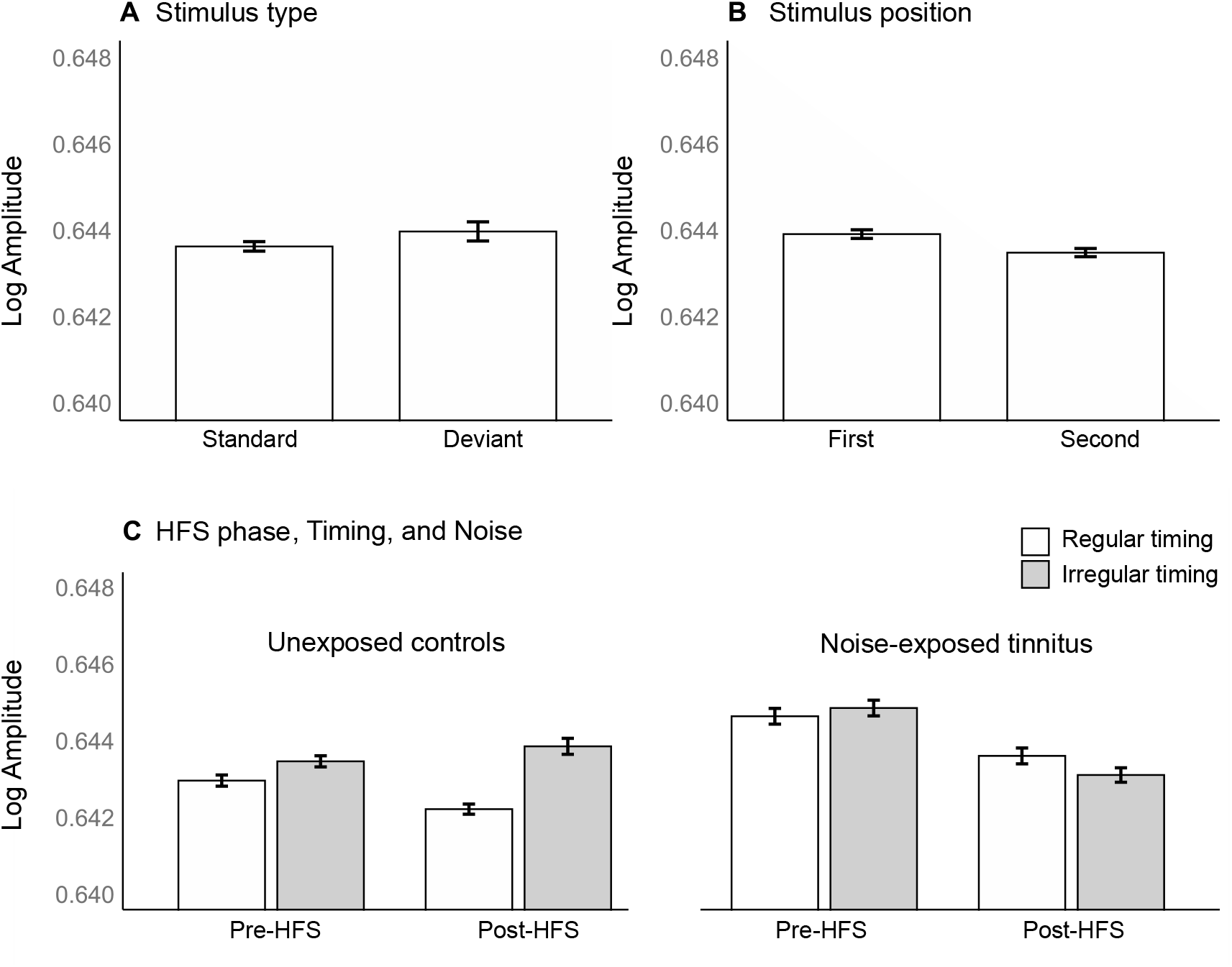
Means (+− standard error of the means) of EP amplitudes demonstrate the main effects of Stimulus type (A), and Stimulus position (B), and the three-way interaction between Timing, HFS Phase, and Noise. Full summary statistics are available in Appendix 1.

Pairwise comparisons investigating the three-way interaction between Noise, Timing, and HFS phase indicated that: only the noise-exposed rats displayed a significant decrease in amplitude from the Pre-HFS phase to the Post-HFS phase in both timing conditions (*ps*<0.001). In contrast, the control group did not show significant changes (*ps*>0.14; Figure 3C). The control group showed significantly higher amplitudes for irregular timing compared to regular timing both pre- and post-HFS (*ps*<0.001). The noise-exposed group, however, did not yield significant differences between timing conditions pre- or post-HFS (*ps*>0.20). These results suggest that timing-related amplitude changes only occurred under normal (control) hearing conditions and were not recovered in the noise-exposed rats following HFS. Finally, amplitudes were significantly higher for the noise-exposed group compared to the control group in both timing conditions in the pre-HFS phase (*ps*<0.007). The noise-exposed group only showed greater amplitudes than controls for the post-HFS phase in the regular timing condition (*p*=0.006) but not the irregular timing condition (*p*=0.47); a one-tailed Bayes factor t test indicated extreme evidence that amplitudes were not higher for noise-exposed rats compared to controls post-HFS in the irregular timing condition (*BF01*=151.4 ± 0.01%). Taken together, these results suggest that noise exposure can impair timing-related SG and that HFS can restore SG to levels similar to that of controls for the unpredictable timing of auditory stimuli.

## Discussion

The current study investigated SG in the MGB of unexposed and noise-exposed tinnitus animal model as well as the effects of HFS. Overall, our findings show the MGB’s distinct role in filtering incoming auditory signals based on the predictability of two stimulus features (pitch and temporal regularity). Further, higher EP amplitudes were found in noise-exposed animals in most conditions (detailed below). Moreover, HFS of the MGB resulted in a decrease in auditory EP amplitudes selectively in noise-exposed animals.

### Auditory filtering capacities of MGB

To our knowledge, this is the first study to provide evidence that auditory SG occurs at the level of the MGB in rats. Responses to deviant tones were significantly enhanced compared to standard tones (Figure 3A). Furthermore, responses to the second tone of a pair were significantly lower than those to the first tone (figure 3B). In control animals, EP amplitudes in the irregular timing condition were significantly higher compared to the regular timing condition (Figure 3C). Here, the higher EP amplitude in response to a deviant tone is comparable to the mismatch responses such as the Mismatch Negativity (MMN) seen in human EEG recordings [30]. A previous study in anesthetized rats showed enhanced EP response to deviant tones across different sleep states (including REM and non-REM sleep) in the auditory cortex [31]. Other studies reported sensitivity of EPs to deviant tones in primary auditory cortex [32]. However, the role of subcortical structures in SG has not yet been elucidated. Overall, the results confirm that the MGB filters sensory information in relation to position, pitch, and temporal regularity.

### Sensory gating after noise exposure

Tinnitus assessment using GPIAS confirmed that noise-exposed animals had tinnitus. Unexposed control rats exhibited higher amplitudes in the MGB for irregular timing compared to regular timing, either before or after HFS. This points to the fact that, in contrast to more popular views of cortical handling of regularity, predictive timing abilities (i.e., entrainment) already occurs at the subcortical level. In contrast, noise-exposed rats were indifferent to timing regularities, either before or after HFS. This may be due to disturbed neurophysiological activity of individual MGB neurons in noise-exposed rats. At a neuronal population level this might be reflected by the inability to distinguish regular and irregular timing in the noise-exposed animals, i.e. to take advantage of the full temporal predictability of the regular stimulus sequence. Thus, at the level of the MGB, noise-exposed animals are unable to distinguish temporal regularity of auditory stimuli. These results suggest that entrainment requires SG and low-level sensory processing, and cannot be restored via MGB HFS. Therefore, findings indicate that entrainment to timing regularities might be a bottom-up process that can be perturbed by disruptions to SG.

Additionally, we found a significantly higher EP amplitude in noise-exposed than control animals across all conditions except for the irregular stimulation group at post-HFS phase. This may be due to the increased synchronization of firing in subcortical auditory centers in noise-exposed animals [33]. As shown before, elevated EPs might be a neural signature of tinnitus [34]. The noise-exposed animals in the current experiment showed tinnitus-like behaviour, which was confirmed by GPIAS. However, prematurely linking elevated EPs in the noise-exposed animal group to tinnitus would be unwarranted. An alternative reason for increased responses may be higher levels of stress in these animals, making them more sensitive to external stimuli. Various studies confirmed alterations in burst patterns in MGB of rats with acoustic trauma which may affect the EP amplitudes of tinnitus animals relative to control animals [34, 35]. Furthermore, a possible confounding effect of hyperacusis cannot be ruled out. Hyperacusis often coexists with tinnitus, and both conditions might be a result of increased synchronized electrophysiological activity, which could lead to elevated auditory EP amplitudes [36]. Follow-up studies are needed in order to disentangle the role of tinnitus, hyperacusis, and hearing loss on disturbed SG. The experimental design should enable to separate animals which show hyperacusis, tinnitus-like behaviour and/or hearing loss or a combination of these. Based on our findings it remains speculative whether the elevated EP amplitudes in the noise-exposed animals were related to tinnitus or hyperacusis.

From a clinical perspective it would be interesting to develop an objective biomarker for tinnitus. The use of SG as an objective biomarker has also been suggested in other disorders in which SG is abnormal. In fact, P50 abnormalities have been found in the EEG of patients suffering from schizophrenia [37, 38]. Although a clinical study suggests that tinnitus follows dysfunctional SG, further studies are needed to confirm this before SG could be considered as a diagnostic tool for tinnitus [10].

### Effect of high-frequency stimulation on sensory gating

We found an overall significant suppression effect of HFS on EP amplitudes in noise-exposed tinnitus animals, but not in noise-unexposed controls. Currently, disrupted signaling as a working mechanism of HFS is the key premise of most hypotheses regarding the restoration of the neuronal physiology in various pathologies [39]. HFS induces action potentials and causes neurons to lose their ability to transmit information [40]. This way, pathological oscillations can be eliminated and neural firing patters can be normalized [41, 42]. The current findings lend preliminary support to this hypothesis; evoked EP hyperactivity, which is a neural correlate of tinnitus, was normalized after HFS, except for stimulus-timing effects. This normalization might be due to a disruption of pathological oscillations in the thalamo-cortical loop, known as thalamo-cortical dysrhythmia [14][4]. This effect was not found in non-exposed control animals, which may indicate the specific influence of HFS on aberrant neuronal firing.

## Conclusions

The MGB serves as a filtering station for auditory stimulus processing. This includes filtering in relation to position, pitch, and temporal regularity. Noise-exposed animals could not distinguish temporal regularity in auditory stimuli. Furthermore, they showed an overall increase in EP activity at MGB relative to unexposed controls. HFS can suppress the increased EP amplitudes towards normal levels and could potentially restore an animal’s sensory gating capacity.

## Abbreviations

DBS: deep brain stimulation
EP: evoked potentials
HFS: high-frequency stimulation
LMEM: linear mixed-effects model
MGB: medial geniculate body
SG: sensory gating

